# Honeycomb crystallography: comb formation under geometric frustrations

**DOI:** 10.1101/2022.03.13.484106

**Authors:** Golnar Gharooni Fard, Daisy Zhang, Francisco López Jiménez, Orit Peleg

**Affiliations:** Department of Computer Science, University of Colorado Boulder, CO, USA; Western Canada High School, Calgary, Canada; Ann and H.J. Smead Aerospace Engineering Sciences, University of Colorado Boulder, CO, USA; BioFrontiers Institute, University of Colorado Boulder, CO, USA; Santa Fe Institute, NM, USA

**Keywords:** honeycomb, topology, pattern formation, self organization

## Abstract

As honeybees build their nests in pre-existing tree cavities, they must deal with dealing with the presence of geometric constraints, resulting in non-regular hexagons and topological defects in the comb. In this work, we study how bees adapt to their environment in order to regulate the comb structure. Specifically, we identify the irregularities in honeycomb structure in the presence of various geometric frustrations. We 3D-print experimental frames with a variety of constraints imposed on the imprinted foundations. The combs constructed by the bees show clear evidence of reoccurring patterns built by bees in response to specific geometric frustrations on these starter frames. Furthermore, using an experimental-modeling framework, we demonstrate that these patterns can be successfully modeled and replicated through a simulated annealing process, in which the minimized potential is a variation of the Lennard-Jones potential that only considers first-neighbor interactions according to a Delaunay triangulation. Our simulation results not only confirm the connection between honeycomb structures and other crystal systems such as graphene, but also show that irregularities in the honeycomb structure can be explained as the result of local interactions between honeybees and their immediate surroundings, leading to emergent global order. Additionally, our computational model can be used to describe specific strategies that bees use to effectively solve each geometric mismatch problem while minimizing cost of comb building.

## 2 Introduction

The wax–made comb of honeybees is a masterpiece of animal architecture, constructed distributively by thousands of bees who create a highly regular hexagonal structure [1]. This storage structure is essential to the survival of the colony and is constructed in a near-optimal minimization of the wax-to-storage space ratio, due to the high energy cost of wax production [2]. In particular, honeybees consume about 8.4 *lb* (3.8 *kg*) of honey to secrete 1 *lb* (454 *g*) of wax [3]. This ratio highlights the importance of the geometry of regular honeycomb [4], as a hexagonal tessellation minimizes boundary-per-area [5–7]. The regular shape of honeycomb cells has intrigued scientists for centuries, from Darwin who postulated that colonies with the least amount of honey waste to create the wax comb structure would be most successful [1], to Thompson who highlighted that honeycombs economize building materials and space [4].

Perhaps even more enigmatic than the regular structure of the comb is the distributed nature of its construction, where worker bees––akin to 3D distributed wax printers–– simultaneously manipulate small pieces of wax to collectively construct a coherent comb structure (Figure 1 A). As honeybees build their nests in pre–existing tree cavities, they deal with situations that do not allow for a regular hexagonal lattice, such as presence of boundaries or structures that require them to combine cells of different sizes, which results in non-regular hexagons and topological defects [8, 9]. Several observations of the structure of the comb [10, 11] suggest that modifications to the regular pattern extend over several cells. This reinforces the hypothesis that a long range awareness of possible constraints results in an adaptive collective behavior with the goal of minimizing the use of wax. While there have been extensive studies focused on the geometry of the hexagonal regular unit cell [12–15], the mechanisms controlling the density and distribution of defects in the lattice are still not well understood. Only in the last decade have there been studies characterizing irregular patterns quantitatively [9, 11, 16], and the rules governing the planning and construction of honeycomb under external constraints incompatible with a regular lattice remain an open question [17, 18].

**Figure 1:**
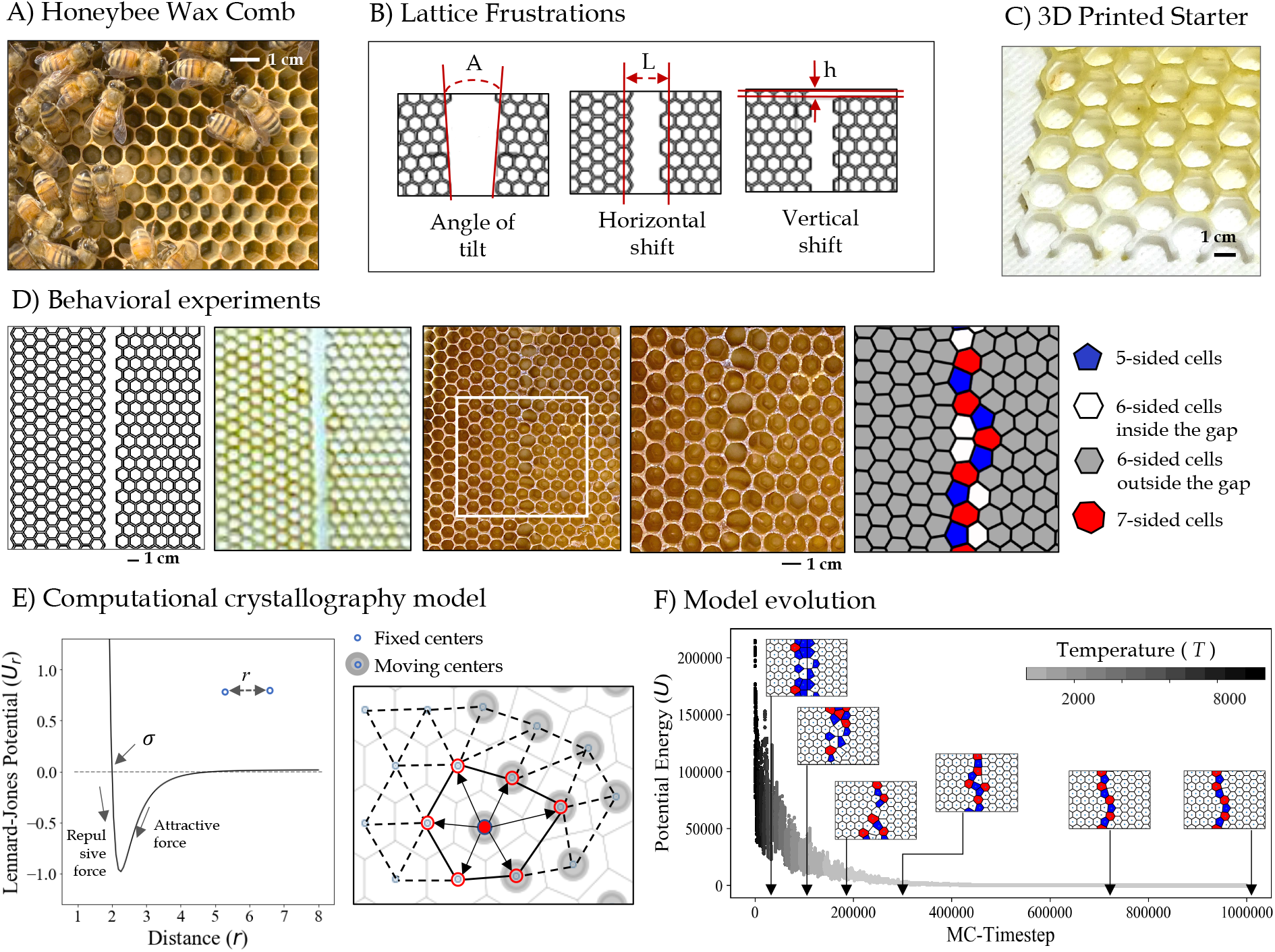
Experiment and model design. A) Collective honeycomb building. B) Illustration of various sources of lattice frustration namely, angle of misalignment (A), horizontal shift (L), and vertical shift (h), introduced through 3D-printed panels. In all these three scenarios there is a *gap* in the given foundation that bees need to fill. C) 3D printed starter frames thinly coated with beeswax. D) Behavioral experiments. From left to right: an example design sheet; the corresponding 3D-printed frame which is coated with a thin layer of wax; the same frame after 25 days when the bees have built comb on it with a white square highlighting the area of interest; the zoomed-in crop of the area of interest; Voronoi reconstruction of the lattice (showing defects in red (*Z* > 6) and blue (*Z* < 6), where the topological charge *Z* is equal to the number of cell sides.) E) Our computational model works according to the rules of the Lenard-Jones potential model. On the left is the graph of the Lennard-Jones potential function: potential energy *U*_*r*_ as a function of the distance *r* of a pair of particles describes both the attractive and repulsive forces between particles. The box on the right shows an illustration of a section of the model with moving and fixed centers highlighted in gray and black. The position of the moving center shown in solid red can be changed at each time step by calculating the potential of its immediate Voronoi neighbors, highlighted with arrows and red rings around them. F) As the model evolves and the system cools within a simulated annealing run, the potential energy decreases. The inset pictures illustrate gradual convergence of the comb structure as the model runs through the optimization process toward the minimum energy state.

In this work, we leverage 3D printing and rapid prototyping to design repeatable experiments with precisely controlled and quantified sources of geometric frustration, illustrated in Fig. 1 B,C. This approach makes it possible to systematically vary a single parameter between different experiments and study its effect on the resulting honeycomb structure. Furthermore, the repeatability enabled by these 3D-printed frames allows us to study statistical variations among experiments with identical initial conditions. We process the resulting patterns using quantitative tools that, while used extensively in crystallography to study lattices in inorganic systems, are not commonly applied to the structures found in animal architecture. In particular, we focus on characterizing the topological defects (*i*.*e*., cells with more or less than six neighbors) that appear as a consequence of the imposed sources of frustration, such as those in Fig. 1 D.

On the modeling front, we also make use of tools from the field of crystallography. We use an algorithm based on simulated annealing to optimize the position of cell centers so that they minimize a variation of the Lenard-Jones potential, which has previously been used to model similar crystallographic structures in graphene [19]. Previous efforts to explain irregularities in honeycomb structure optimized the geometry within a fixed initial topology of the cell lattice [11], which could effectively restrict the configuration space. In contrast, our model optimizes the geometry and topology of the lattices simultaneously by allowing the interconnections between the cells to change and adapt to the nearest neighbors’ potential. We assume the bees are implicit. Conceptually, the bees would position the cells according to the local interactions with adjacent cells, as defined in the Section 3. The boundary conditions in our algorithm closely replicate the experiments, allowing us to quantitatively compare experimental results with model predictions. At the end of the simulation, the arrangement of the cells reach an equilibrium configuration that can be directly compared against the configurations bees create in our experiments.

## 3 Methods

We take a perturbation approach to reverse-engineer the local behavioral rules that lead to honeycomb construction: by perturbing the system with carefully prescribed conditions, we anticipate gaining a deeper understanding of the underlying principles of distributive construction. Our goal is to understand how bees overcome scenarios in which engineered constraints make it impossible to build a regular hexagonal lattice. Our focus is on three different cases of frustrations, namely angle of tilt (*A*), horizontal shift (*L*), and vertical shift (*h*) (illustrated on sections from our design sheets in Fig. 1B). Both the behavioral experiments and our computational model are designed in a way that enable us to vary the values of these three parameters and study their impact on the honeycomb lattice.

### 3.1 Behavioral experiments

We introduce geometric frustration to the system on a microscopic scale (*i*.*e*. on the scale of individual cells) using 3D-printed foundation plates. The printed foundation is only introduced to segments of the plate, and the geometry and patterning of the panels is deliberately designed so that these regions cannot be connected using a regular hexagonal honeycomb. For example, the *gaps* within the printed patterns can be designed so that the bees will not be able to simply extend the provided foundation. Fig. 1 D shows the result of our behavioral experiments, starting with a sample of a design sheet followed by the corresponding 3D-printed plate, which is provided to the bees as a starter frame. We then take pictures of the fully-built frames and identify areas of interest on each image. We define this area as the largest crop on each plate that contains an undamaged comb with cells within and on either side of the gaps, highlighted with a white rectangle in the middle image in Fig. 1 D. We use computer vision techniques to automatically identify individual bee comb cells on the selected crop. For details and examples of the steps in this process please refer to Fig. S2. The final image in the series in Fig. 1 D is an example of the output of the image processing procedure: a Voronoi tiling based on the cell centers, where each comb cell is replaced by the corresponding Voronoi cell which reveals the non-regularity of the shapes of cells within and around gaps. We find striking agreement between the Voronoi construction and the network of honeycomb cells built on the experimental frames under various conditions. Nevertheless, to confirm that the Voronoi reconstruction matches the actual comb image, we visually inspect each image with the Voronoi diagram overlaid and make corrections if necessary. Once the corrected Voronoi tessellation is created, it can be used to calculate geometric and topological properties of the cells such as coordinates of the cell center, topological charge *Z* (*i*.*e*., number of cell neighbors), and cell area.

### 3.2 Computational crystallography model

Our goal in this section is to develop a physics-based mathematical description of a set of rules governing honeycomb construction at the local scale, that can explain the global patterns we observe in our experiments. To that end, we establish a computational model solved via simulated annealing, a technique for approximating the global optimum of a function [20, 21] that is based on Monte Carlo methods and was originally developed to generate sample states of a thermodynamic system [22]. It receives its name from the similarity to the process of annealing in materials science and is often used to study the formation of crystals resulting from the minimization of a potential energy [23–25] or to reconstruct the microstructure of dispersions and heterogeneous solids [26–28]. It requires defining a function to be minimized (analogous to the internal energy in a crystal), which depends on the state of the system. In our case, state variables are the position of the centers of honeycomb cells, which are used to define a potential energy function *U*. The method minimizes the potential by exploring neighbors of the current state of the problem, in which particles are disturbed by a small displacement. Each displacement is accepted according to a probability *P* given by:

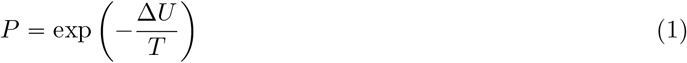

where Δ*U* is the change in potential energy due to the change from the current to the new proposed state, and *T* is a global parameter that controls the probability of accepting changes in state that increase the energy, and is analogous to temperature in real physical material annealing. Temperature is traditionally decreased during the process. It is initially high, so that the initial exploration of the solution space accepts a wide range of possible states, including some that increase the energy. The temperature then decreases as the solution converges to an optimum, so that only changes that minimize the energy are accepted, as in Fig. 1F. The specific range of values depends on the problem and is often unrelated to realistic temperature values in metallurgy processes.

The minimized potential is a variation of the Lennard-Jones potential (Fig. 1 E), known to produce hexagonal lattices in the absence of constraints. The interaction between particles is given by:

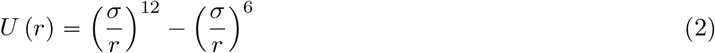

where *r* is the distance between two particles that are identified as first neighbors by a Delaunay triangulation (*i*.*e*., we ignore long-range interactions), and *σ* is the distance at which the potential is zero. The potential is minimized at a particle-to-particle distance *d*, given by *d* = 2^1/6^ *σ*. In our case *d* = 5.40 mm, measured directly from regular hexagons in control experiments. The Delaunay triangulation is updated at each step, to account for the interplay between geometry and topology in our network as the the distance between particles change.

The model imposes the same constraints used in the experiments: misalignment angles, *A*, and horizontal and vertical shift of the hexagonal lattices, *L, h*. The algorithm considers two types of particles: the center of the cells imprinted in the starter panel, which act as a set boundary (*i*.*e*. fixed centers, shown with a blue dot in Fig. 1 E), and the center of the cells that are created in the gaps, which are the variables in the minimization process (i.e. moving centers, highlighted with a gray shade around them in Fig. 1 E). The initial arrangements of the fixed cells in the model are chosen to replicate the scenarios explored with our experiments. The number of moving centers in the simulations depends on the size of the system, as well as the specific type of geometric frustration being explored, and is based on the number of cells created by the bees in our experiments (see Fig. S9 for a complete list of these values derived from the experiments for each parameter combination).

## 4 Results

In this section, we present our experimental and modeling results, focusing on the effect of varying each of the sources of geometric frustration, namely misalignment angles (*A*), horizontal shift (*L*), and vertical shift (*h*) of the regular lattices. The horizontal shift is expressed as a function of *d*, the distance between the centers of two adjacent hexagons in the regular lattice, which is used to define our simu<u>l</u>ation framework, as described in Section 3.2. The vertical shift will be expressed as a function of 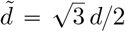, the vertical distance between two adjacent cells at 30° from each other. The results that follow use data collected from a total of ten A. *mellifera* honeybee hives that built comb on the 3D-printed experimental frames with varying values for the three parameters described above. The crops that show the qualitative agreement between the model and experiments are presented in panel A of Fig. 2-Fig. 4. These results display the most common patterns that we observed in both simulations and experimental data under each condition. A quantitative analysis of these results is performed on a larger dataset containing all of the experimental data (3-5 crops of the same size for each parameter combination) and 10 simulation runs. The largest common crop size across all of the experimental data displayed in this work contains 22 rows of cells. To generate model results, the size of the simulation box is set to match the experimental crops for better visual comparison. All plots show the average of all identical experiments and simulations, with the standard deviation as error bars. Finally, in order to verify that the resulting patterns are not affected by external factors (*e*.*g*. material used in 3D-printing), and arise from the geometric frustrations that were imposed on the starter frames, we 3D-print several control frames without any geometric frustrations (see Fig. S1 for instances of experimental and control frames), and place them in all of the hives along with the experimental frames. Unsurprisingly, we find that the bees consistently build regular and perfect comb on the control frames without any defects. Please see Fig. S3 for the results of running the image processing pipeline and automatic cell detection on an instance of a control frame.

**Figure 2:**
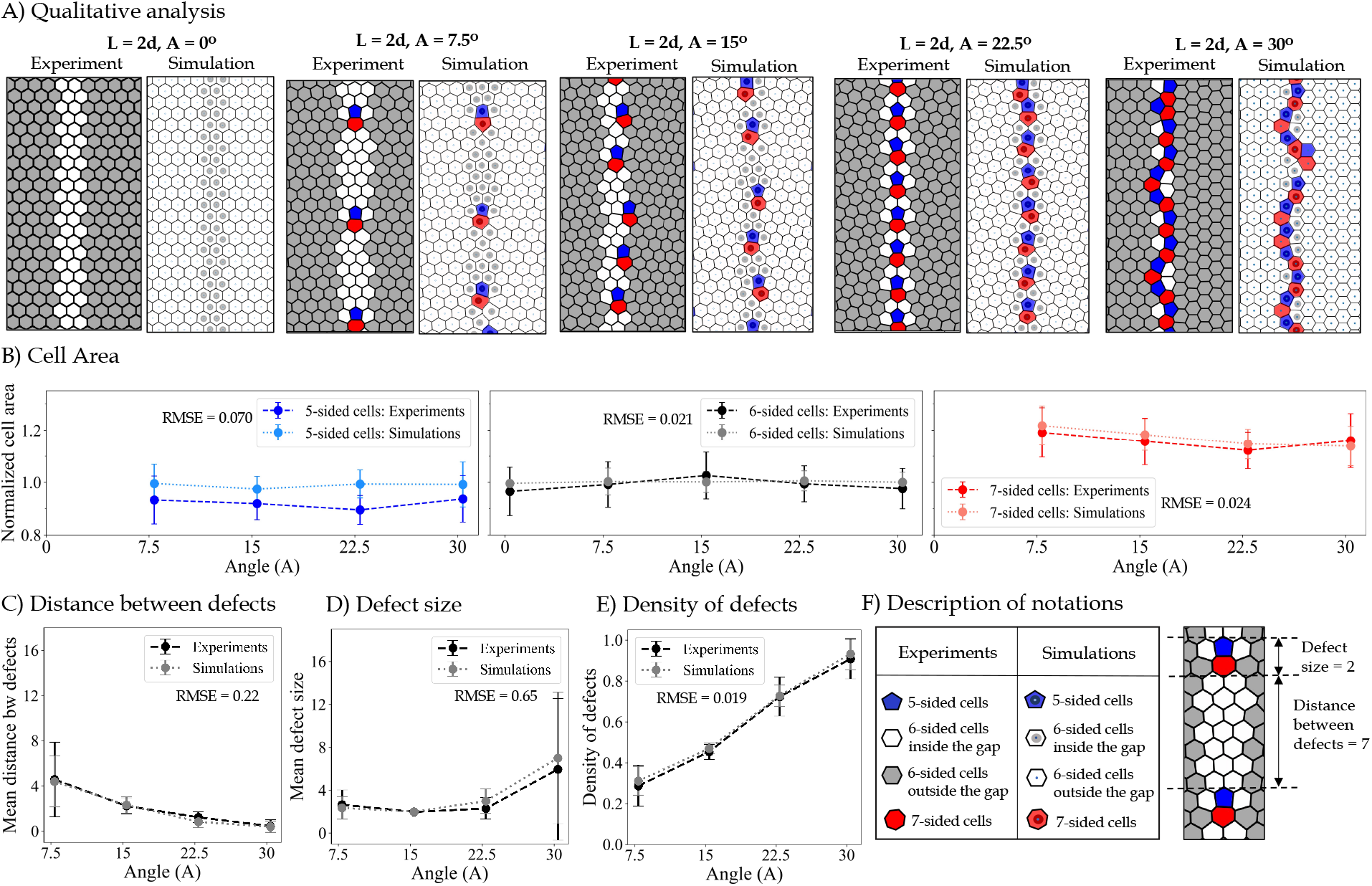
Results for varying the misalignment angles (0° ⩽ *A* ⩽30°). A) Pairs of images taken from experiments (on the left) and simulations (on the right) to highlight the qualitative agreement between our model and experiments. Defects are non-hexagonal cells, shown in red (*Z* = 6) and blue (*Z* = 6), where *Z* is the topological charge (*i*.*e*., number of sides) of each cell. B) Mean cell area across different angles categorized by the cell shape, shown both in experiments (dashed lines with darker colors) and model (dotted lines and brighter colors). C) Mean distance between defects in experiments (dashed black) and simulations (dotted gray) shows a decline as the angle of misalignment increases. D) Average length of defect chains in experiments (dashed black) and simulations (dotted gray) generally increases as the angle of misalignment increases. The large error bars in the case of *A* = 30 suggest the presence of defective chains of various sizes when the angle of misalignment is large. E) Density of defects (*i*.*e*. the number of non-hexagonal cells divided by the height of the crop) shows a sharp increase as the angle of misalignment increases. To quantify the agreement between experiments and model results, the value of root mean squared error (RMSE) is calculated and shown on for all the parameters plotted in panels B-E. The small values of RMSE indicate that our model can predict the experimental data relatively accurately. F) The table on the left shows a description of the notations used in panel A to distinguish between various cell types. On the right is an example of how the two variables in panel C and D are calculated.

### 4.1 Exploring the impact of the angle of misalignment (A)

Fig. 2 A shows the qualitative comparison between the processed experimental frames and the simulation output with various values of tilt. In order to focus on the impact of specific angles in the pattern of the cells that connect the gap between the panels, Fig. 2 A reports instances of both experimental and model results with varying angle (0° ⩽ *A* ⩽ 30°) but with fixed distance of *L* = 2*d* between the panels. We show five pairs of images in Fig. 2 A, with Voronoi reconstruction of our experimental data on the left. The shades of gray in the experimental results represent the cells built on the given 3D-printed foundations whereas the cells built inside the gap with no foundation are shown as white. In the model results, shown on the right hand side of the pairs in Fig. 2 A, the moving lattice elements (equivalent to cells built in the gap) can be distinguished by darker centers. Both the model and the experiments resolve the frustration through the introduction of topological defects (i.e., cells with other than six neighbors). These often take the form of dislocations, *i*.*e*. a duplet of a positive and a negative (5-7) defect.

The results in Fig. 2 A show that increasing the misalignment angle increases the linear density of defects (*i*.*e*., decreases the distance between pairs), with excellent quantitative agreement between the model and the experiments. We also explore the variation in cell sizes by calculating cell areas across all angles, and categorizing the results based on the number of sides, as shown in Fig. 2 B. Unsurprisingly, cells with a greater number of sides (shown in red) are on average larger. This pattern persists across different angles of tilt both in experiments and simulations. However, the distribution of cell sizes shown in Fig. S4 denotes that there is more variability of the cell sizes in our experimental data compared to the cell sizes generated in the model. This can be due to specific functionality of cells of various sizes and shapes within the hive. For instance, we find that many of the larger hexagons and heptagons that emerge as a result of tilted patterns on our experimental frames are used as drone comb (see Fig. S8), which typically have larger cells for the queen to lay unfertilized eggs in. Fig. 2 C captures the opposite correlation between increasing the angle (*A*) and the distance between the defects. The distance between defects is calculated by counting the topological distance between two sets of defective cells, see Fig. 2 F for an example of a distance of seven cells between two dislocations. Furthermore, we plot the change in the average size of defect chains as a function of angle of tilt in Fig. 2 D, which shows a positive correlation between increasing the angle of tilt and having larger defective sequences. The size of a defect chain is calculated by counting the number of connected (uninterrupted) non-hexagonal cells. See Fig. 2 F for an example of a defect size of 2. Fig. 2 E highlights the positive correlation between the number of defective cells per crop and the angle of tilt. Since there are no defects in the case of *A* = 0°, this angle is not included in the statistical analysis of the defects

### 4.2 Exploring the impact of horizontal shift (L)

Fig. 3 A illustrates how the structure of the comb is impacted by keeping a fixed angle of tilt *A* = 30° across all instances while increasing the distance between them in range 2*d* ⩽ *L* ⩽ 4*d*. We also perform experiments in which the distance between the given hexagonal lattices goes up to 11*d*. However, during the time frame of our experiments, the bees did not connect the two lattices if the distance between them is larger than *L* = 4*d*. See Fig. S7 for an example of a comb built on frames with large gaps between the given regions. In fact, even within the range of 2*d* ⩽ *L* ⩽ 4*d*, the size of our experimental dataset shrinks as we increase the distance, with fewer analyzable samples in *L* = 4*d*. As Fig. 3 A illustrates, bees use long chains of alternating 5-7-sided cells inside the gaps across various distances when the angle of tilt is large (*A* = 30°). When the distance between the panels is larger than *L* = 2*d*, the bees continue to build hexagons even when there is no foundation underneath and then use a mix of 5-7-sided cells to connect the hexagonal lattices on either side of the gap.

**Figure 3:**
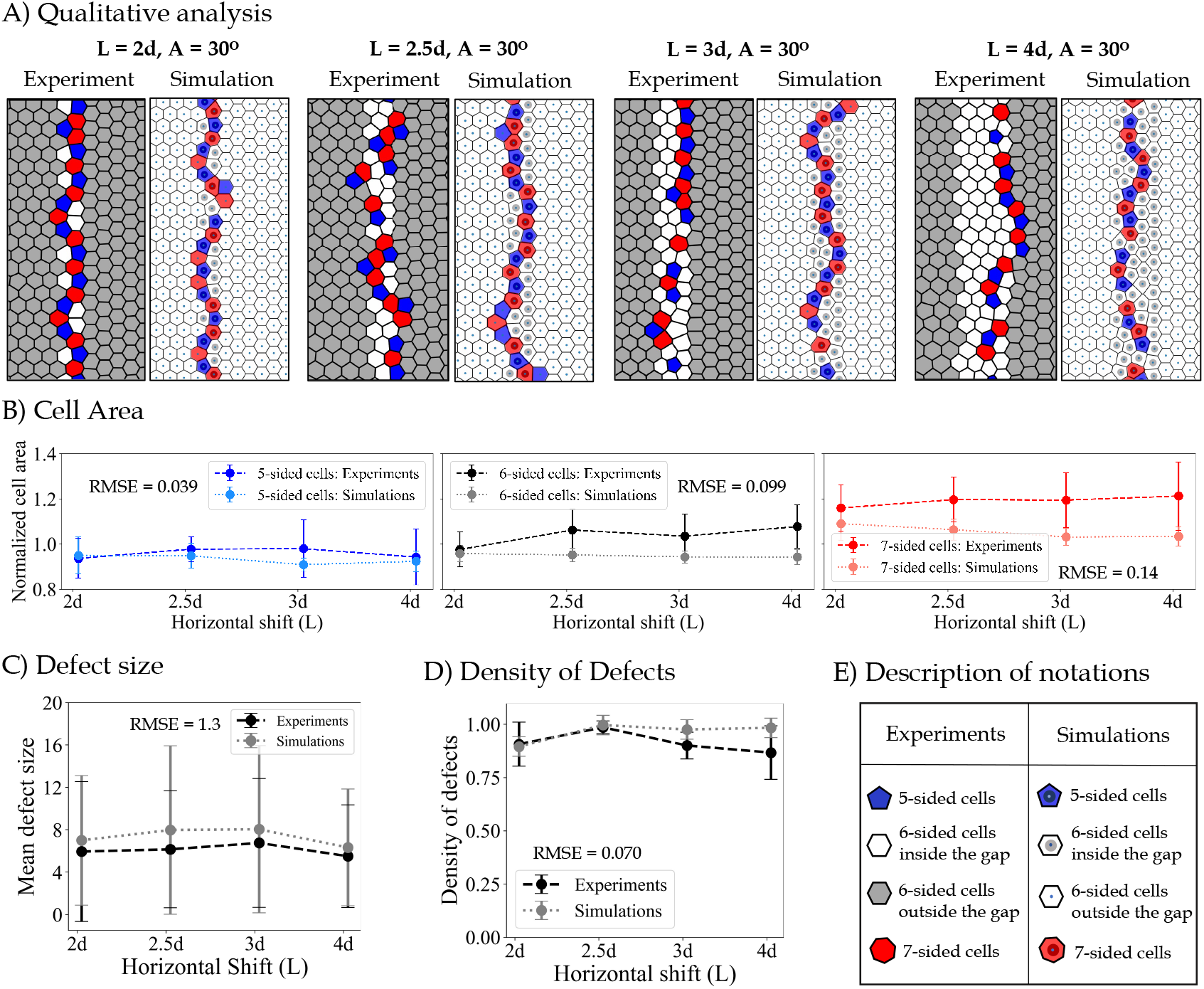
Results for varying the horizontal shift parameter 2*d* ⩽ *L* ⩽ 4*d*. A) Pairs of images taken from experiments (on the left) and simulations (on the right) to highlight the qualitative agreement between our model and experimental results. Defects are shown in red (*Z* > 6) and blue (*Z* < 6), where *Z* is the topological charge (*i*.*e*., number of sides) of each cell. B) Mean cell area across different values of vertical shift categorized by the cell shape, shown both for experiments (dashed lines with darker colors) and model (dotted lines and brighter colors). To see the distribution of cell sizes refer to Fig. S5. C) Average length of defect chains in experiments (dashed black) and simulations (dotted gray) shows small changes for various horizontal distances between panels. The large error bars indicate the presence of defective chains of various sizes when the angle of misalignment is *A* = 30. D) Density of defects (*i*.*e*. the number of non-hexagonal cells divided by the height of the crop) shows a high density of defective cells within the gap. The density of defects stays relatively high as the value of the horizontal shift increases. E) The description of the notations used in panel A. shown in Fig. 2 C-E.

Comparison of cell sizes shown in Fig. 3 B, confirms the positive correlation between the number of sides and cell area which is quantified in the previous section as well. This pattern holds the same across various distances, both in the experiments and in the model results. Fig. 3 C indicates that the average size of defects does not change substantially across varying distances. This is mainly because the angle of tilt is constant (*A* = 30°) so the bees continue to build their preferred shape (hexagons) within the gap (regardless of the horizontal distance between the panels) and finally use the long chains of defective cells to combine the two sections of comb. The relatively high but constant density of defects, shown in Fig. 3 D, confirms the qualitative results shown in panel A, suggesting that horizontal shift between panels does not impact the size or the density of defects as long as the angle of misalignment is fixed. In fact, our experimental data (see Fig. S6) confirms that bees build the same patterns of defective cells, shown in Fig. 2 A, to combine the two lattices of various angles of misalignment even when the distance between the two lattices is larger than *L* = 2*d*.

### 4.3 Exploring the impact of vertical shift (*h*)

In this section, we explore the impact of imposing a vertical shift, *h*, to the geometry of two panels at either side of the gaps. To highlight the isolated effect of *h*, we illustrate two conditions, namely *h* = 0 where there is no vertical displacement of the panels, and 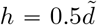, while assuming a fixed value of *A* = 0°, and *L* = 2*d*, so that the vertical shift is the only source of frustration. Our qualitative results for these two scenarios are shown in Fig. 4 A, which demonstrates great agreement between experiments and simulations showing more defective cells used for filling the gap between the given hexagonal regions when there is a vertical shift in the structure of the panels. As shown before, there are no defects built in the gap when *L* = 2*d, A* = 0°, and *h* = 0. However, the pairs (or short chains) of defective cells are built inside the gap as a result of a non-zero value of *h*. We also notice the common tilt of these often short chains of defects and quantify it across both experiments and simulations. Fig. 4 B shows two distributions for the angle of defects across experiments (solid bars) and simulations (hatched bars) with a very similar mean value highlighted with a dashed red line on both plots (*i*.*e. α* = 24.97° in the experiments and *α* = 27.14° in simulations). The average cell size plots shown in Fig. 4 C, follow the same trend as described before, where the cells with fewer sides are on average smaller. Since there are no defects in case of *A* = 0, *h* = 0, only six-sided cells are shown in these size plots. The average distance between defects, defect size, and density for *A* = 0, *h* = 0.5*d* are shown in Fig. 4 E-G for experiments and simulations. The large value of the average distance between defects shown in Fig. 4 E suggests sporadic, small chains of defects, when *h* = 0.5*d*, which is confirmed in Fig. 4 F-G. Interestingly, even if the distortion created by vertical shift could be resolved with deformation of the hexagonal cells without introducing topological defects, we observe similar non-zero defect density in both experiments and simulations. It is not clear if the presence of defects is used to reduce distortion of the hexagonal cells or to create a variation in cell size for different functionality.

**Figure 4:**
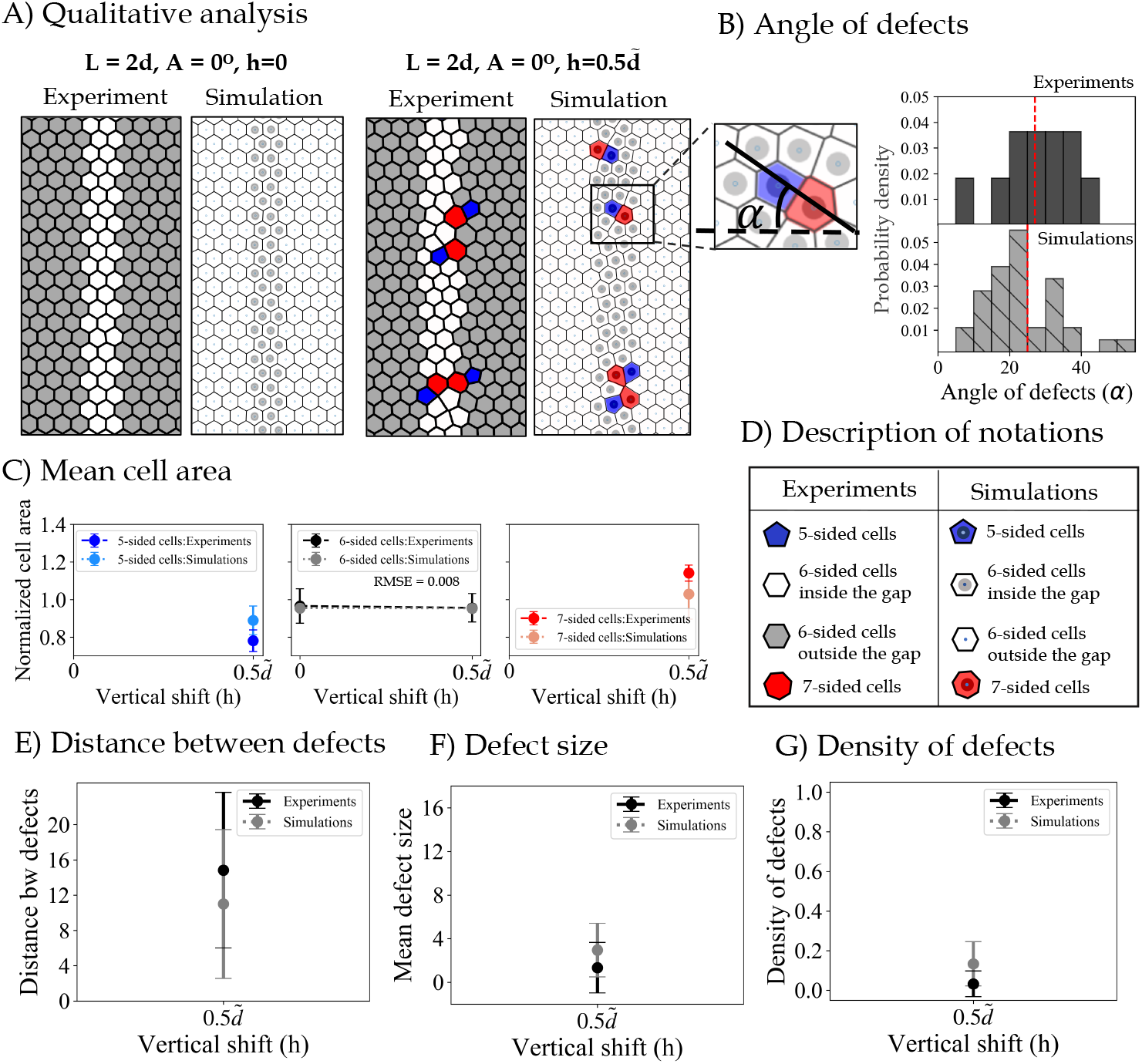
Results showing the impact of vertical shift *h* = 0, 0.5*d*. A) Pairs of images taken from experiments (on the left) and simulations (on the right) which highlight the qualitative agreement between our model and experimental results (showing defects in red (*Z* > 6) and blue (*Z* < 6), where *Z* is the number of cell sides). B) Probability density of the angle of defects *α* shows very similar mean and standard deviation values across both experiments and simulations. *α* = 24.97° ± 10.03° in the experiments and *α* = 27.14° ± 11.05° in simulations, and 0° < *α* < 90°. The mean values are highlighted with dashed red lines on both plots. C) mean cell area categorized by the cell type, shown in experiments and model for the two values of *h*. D) Description of the notations used for various cell types shown in panel A. E) Mean distance between defects in experiments (dashed black) and simulations (dotted gray). F) Average size of connected defective cells in experiments (dashed black) matches with our simulations (dotted gray) G) Density of defects (*i*.*e*. the number of non-hexagonal cells divided by the height of the crop).

## 5 Discussion

Several processes in nature result in the formation of self-organized lattice patterns, including graphene at the nano-scale [29,30], colloidal crystals at the micro-scale [31, 32], and elastic dimples in soft bi-layers at the macro-scale [33,34],. In most cases, the process can be explained as the result of minimizing a potential given by the interactions of the particle, such as inter-molecular potential between two atoms [35] or electromagnetic potential between charged particles [36]. In the absence of external constraints, the particle distribution that minimizes the potential is a topologically and geometrically regular crystal lattice [37]. However, defects such as dislocations or disclinations can appear due to different sources of geometric frustration, such as incompatibilities between the lattice and its boundary or between two crystalline regions with different orientations [38–44].

Inspired by the similarities between the grain boundaries in our system and those observed in graphene [45], we have used a variation of Lennard-Jones potential in our modeling of the honeycomb formation under geometric frustrations. The agreement between simulations and experiments in our study further reinforces previous observations that defect formation is relatively insensitive to the type of interaction between lattice elements, so that similar structures are observed across different systems [46, 47]. Different aspects of comb construction might be the result of global planning or local interactions. When it comes to irregularities within the honeycomb pattern, our model reproduces them based on local interactions alone.

While our work was in progress, we became aware of a related study by Smith et al. [11]. Our conclusions are in agreement with respect to the 5- and 7-sided cells repeated pattern, which was observed in the absence of a frame foundation. In addition, we have shown that using 3D foundations to introduce geometrical frustration makes it possible to systematically vary a single parameter across different experiments and rationalize its effect on the resulting honeycomb structure. Furthermore, the repeatability provided using these 3D-printed frames allows us to study statistical variations among experiments with identical initial conditions. Lastly, our model optimizes the geometry and topology of the lattices simultaneously by allowing the interconnections between the cells to change and adapt to the nearest neighbors’ potential. The boundary conditions in our algorithm closely replicate the frustrations imposed in experiments, allowing us to directly compare experimental results with model predictions quantitatively.

Our combined experimental and modeling framework can be expanded to directly address more specific hypotheses regarding the optimization process that the bees perform when constructing the honeycomb. In particular, instead of using atomic-based variations, the potential minimized through simulated annealing could be constructed based on lattice parameters that are assumed to be of importance to the bees (*e*.*g*., uniform size and aspect ratio). The combination of experiments exploring a wider range of sources of frustration (*e*.*g*. boundaries, curvature) and a systematic exploration of potentials will enable a deeper understanding of the decision-making mechanism behind the bees’ choices of constructing cells of various shapes and sizes. Conversely, by understanding honeybee comb construction –honed by evolution, selection, and refinement––we can expect to develop new rules for bio-inspired structural design of cellular solids. These are of particular interest for 3D-printing and other advanced manufacturing techniques, which are not limited to using fully periodic lattices.

## 6 Acknowledgements

O.P. and F.L.J acknowledge funding from the University of Colorado Boulder RIO Seed Grant Program. We also acknowledge funding from the University of Colorado Boulder, BioFrontiers Institute (internal funds). We thank Seneca Kristjonsdottir and Christopher Borke for bee management, Paul Bontempo and Ashley Atkins for their assistance with data collection and organization, Riley Perez for designing the 3D-printed frames and Olga Shishkov for reading and commenting on the manuscript. We thank Prof. Elizabeth Bradley, Prof. Greg Stephens, and members of the O.P. laboratory for insightful feedback and discussions.

