## Supplementary Material for "Honeycomb crystallography: comb formation under geometric frustrations"

### 1 Supplementary Information

#### 1.1 Experimental Design

To quantify the comb construction, we establish a behavioral assay and a high–throughput analysis method to track the constructed comb on each image. Parameters under study are varied systematically, so that their effect can be isolated and quantified. These careful measurements allow us to assess the construction dynamics and final configuration of the honeycomb. All the design work for this study is conducted using the SolidWorks 2019 CAD design software [1]. All of our experiments are performed using colonies of the European honeybees *Apis melifera* L. at the Peleg lab apiary in Boulder, Colorado. To collect data, we use langstroth hives with 19 $in$ (480 $mm$) deep frames, with $9\frac{1}{8}$ $in$ (230 $mm$) depth, and $1\frac{3}{8}$ $in$ (35 $mm$) width, see Fig. S1. Each frame consists of a pair of 3D printed plates of the size $212.00 \times 215.90$ $mm$, that are placed opposite to each other. Each hexagonal element on a 3D-printed plates has a side-length of 3.12 $mm$, which is obtained by averaging the cell size of naturally occurring honeycomb. To account for at least three repetitions of all the combinations of parameters under study, We scatter a total of 50 3D-printed experimental frames into ten honeybee colonies and fill the rest of the space in the hives with normal frames with full plastic foundations.

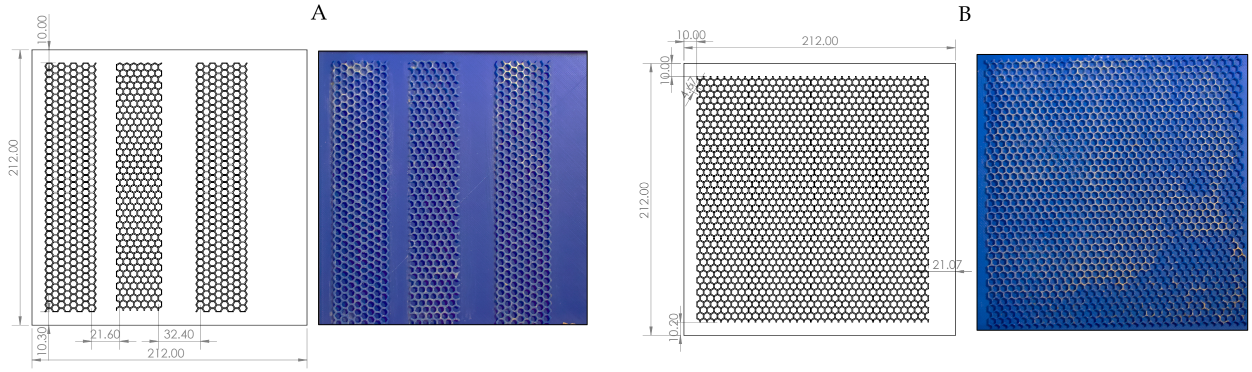


Fig. S1: **Experimental v. control plates.** A) An instance of an experimental design sheet (on the left) with the corresponding 3D-printed plate covered with beeswax (on the right). Each experimental design sheet contains two or three geometrical frustrations with a unique parameter combination. The number of experimental plates is set so that each parameter combination is replicated 5 times. B) A control frame design sheet with no geometric frustrations (on the left) and the corresponding 3D-printed plate covered with beeswax (on the right).

To promote comb building we coated the printed foundation with a very thin layer of natural beeswax. The progress of comb building is regularly checked by taking pictures of both sides of all our experimental frames, twice a week. Given that the speed of comb building is not the same across all of the hives, these regular photography sessions ensures that we do not miss the fully-built stage of the comb before the cells are capped with honey or brood. To minimize the disturbance caused by the recurring photography sessions, we build our photography setup inside the apiary so that we can instantly return all the frames back to the same position we took them from inside the hives. Once the experimental frames are fully built, they are taken out of the hives and we bring them inside the lab for safekeeping. On average, it took about two months for all the 50 frames to be complete. We obtain high-quality images of the comb using a Nikon DSLR camera and controlled lighting, and a black background. The camera was mounted on a fixed metal structure $2m$ from the comb to avoid distortion at the edges of the image and to keep a constant pixel-to-millimeter relationship. At this distance and camera resolution, 1 $mm =$ 15 pixels.

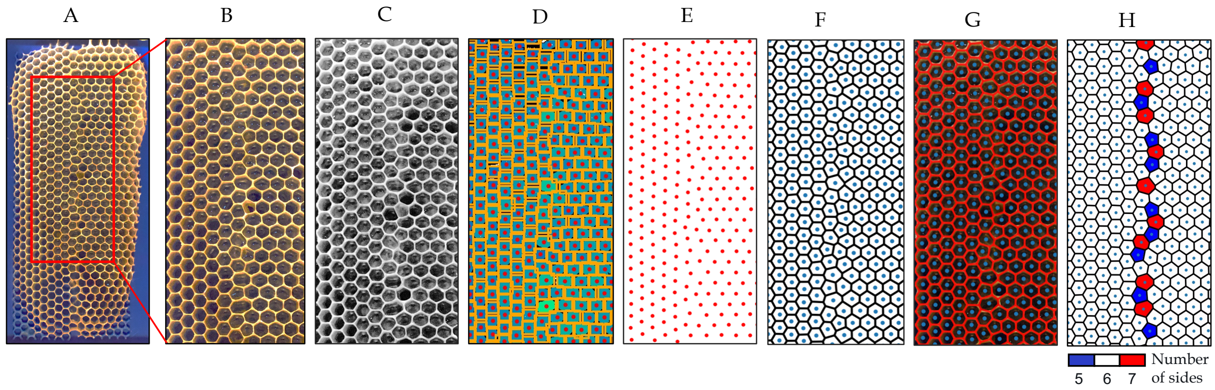


Fig. S2: Experimental analysis pipeline described on a sample 3D-printed frame with $L = 2d$, $A = 30°$, and $h = 0$. A) Comb on the experimental frame with a specified region of interest B) cropped image C) contrast correction by histogram equalization D) automatic cell detection after thresholding and segmentation. Individual cells are highlighted by orange rectangles and red centers. E) Point cloud extraction from the cell centers. F) Voronoi tessellation G) Overlay of the Voronoi reconstruction on the initial image in B. H) Coloring Voronoi cells by the number of neighbors/sides, brings out the defects used for filling the gap between two hexagonal panels with misalignment of $A = 30°$.

#### 1.2 Image Processing Pipeline

Our image processing pipeline starts with focusing on the region of interest highlighted with a red rectangle in Fig. S2 A, which is cropped in Fig. S2 B. We then perform a simple histogram equalization on the selected region to improve the contrast of the image, Fig. S2 C. Segmentation is done using the *felzenszwalb* algorithm [2] which marks the boundaries of the cells. We then used Otsu's method of thresholding [3] along with a few morphological filters for detecting individual cells and their centers, shown in Fig. S2 D. From the position of the centers of these cells, we generate a point cloud, shown in Fig. S2 E, and use it to construct a Voronoi tessellation in Fig. S2 F. To validate the output of the automatic process of detecting cells and reconstructing the comb structure, we overlay the Voronoi tessellation output on the initial crop in Fig. S2 G and manually fix any discrepancies. Finally, we color the cells based on their number of sides, shown in, Fig. S2 H. Additionally, we run the same image processing steps on all of the control frames. The outputs of every step is shown in Fig. S3 A-G. The pdf of the cell size distribution shown in Fig. S3 H shows that in the absence of geometric frustrations bees only build regular hexagonal cells.

#### 1.3 Distribution of Cell Areas

Fig. S4 shows the histogram of cell sizes for the configurations depicted in Fig. **??** A, for both experiments (top row) and simulations (bottom row). The value of horizontal shift is fixed at $L = 2d$ and the value of angle of misalignment is in the range $[0, 30]$, and each column shows the probability density of each cell type highlighted with different colors. The probability density of different cells is categorized by the number of sides (*i.e.* 5,6, and 7). Overall, a couple of 4-sided and four 8-sided cells appeared in both experimental and model analysis but we did not include them. Note that when the misalignment angle $A = 0°$, both bees and the model have no trouble filling the horizontal distance of $L = 2d$ with hexagons and the non-hexagonal cells emerge as a result of higher angles of misalignment between the given hexagonal panels. Similarly, Fig. S5 shows the histogram of cell sizes for the configurations depicted in Fig. 4, for both experiments (top row) and simulations (bottom row). The value of the misalignment angle is fixed at $A = 30°$ and the value of horizontal shift is in the range $[2d - 4d]$, and each column shows the probability density of each cell type highlighted with different colors. The distribution of the cell sizes created by the bees on the experimental frames, shown in the top rows of Fig. S4 and Fig. S5, are generally wider when compared to the cell sizes generated using the model. This suggests that the model is producing cells with relatively similar sizes.

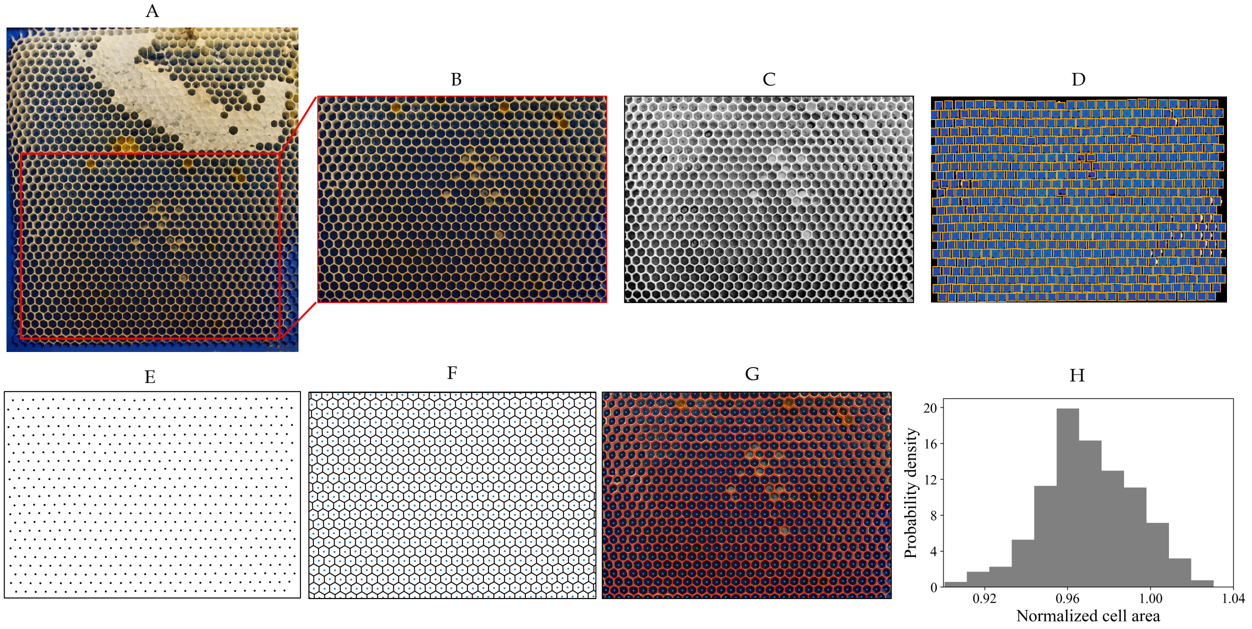


Fig. S3: Image processing analysis performed on an example control frame with no geometrical frustrations on the foundations printed on the starter frame. A) Comb built on the control frame with a a specified region of interest B) cropped image C) contrast correction by histogram equalization D) automatic cell detection after thresholding and segmentation. Individual cells are highlighted by orange rectangles and red centers. E) Point cloud extraction from the cell centers. F) Voronoi tessellation G) Overlay of the Voronoi reconstruction on the initial image in panel B. H) Probability density of the cell areas in panel B, normalized by the size of a perfect hexagon, shows that the bees follow the regular patterns on the foundations given on the starter frames and hexagonal cells are built in the absence of geometric frustrations.

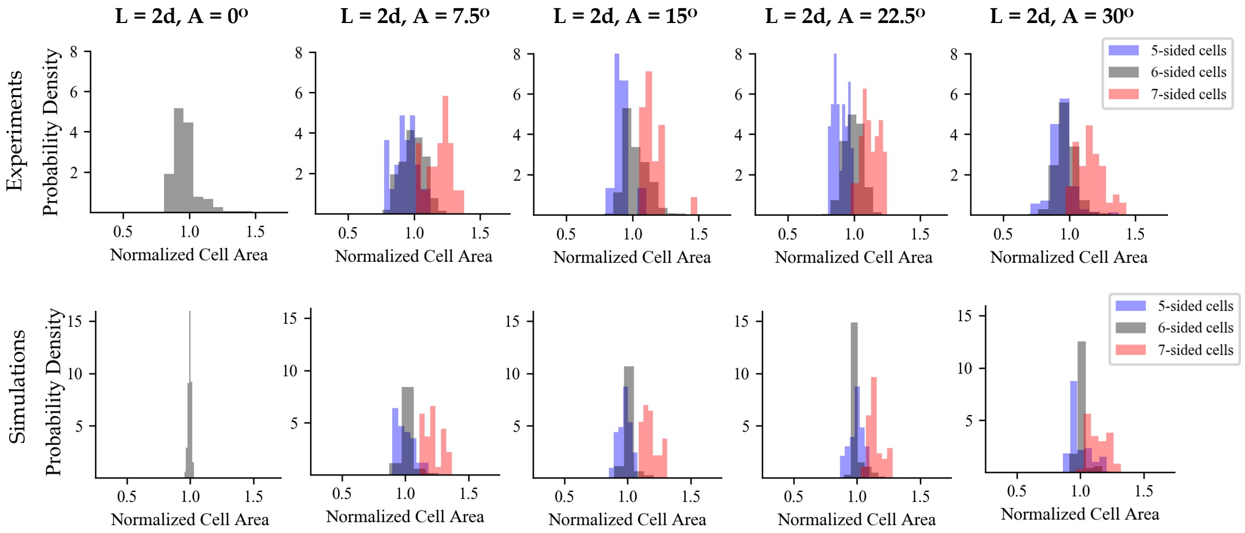


Fig. S4: Probability distribution of cell sizes for every angle in experiments (top row) and simulations (bottom row).

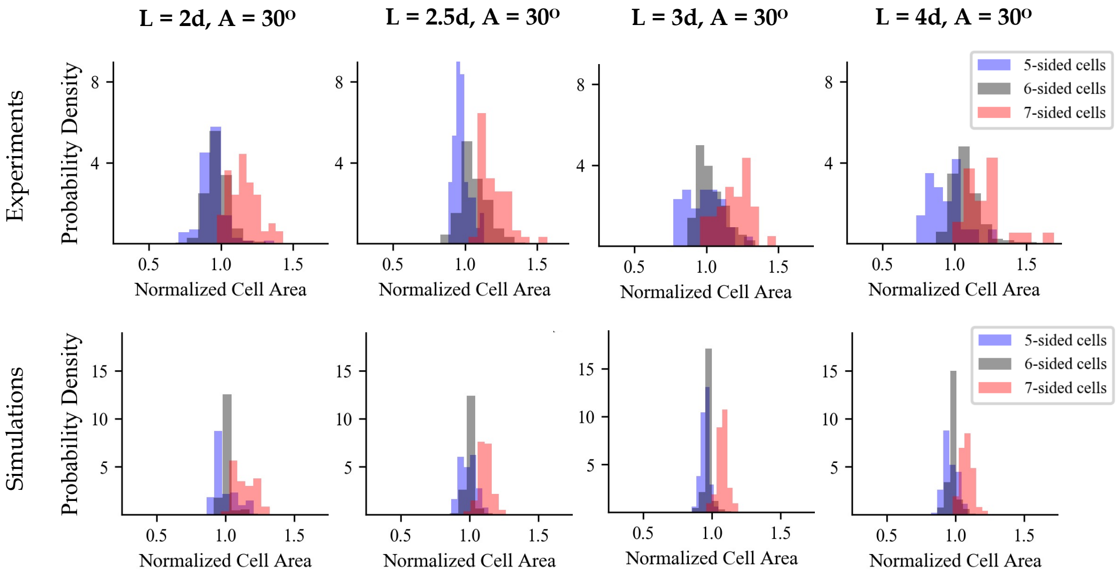


Fig. S5: Probability distribution of cell sizes for all the values of horizontal shift in experiments (top row) and simulations (bottom row).

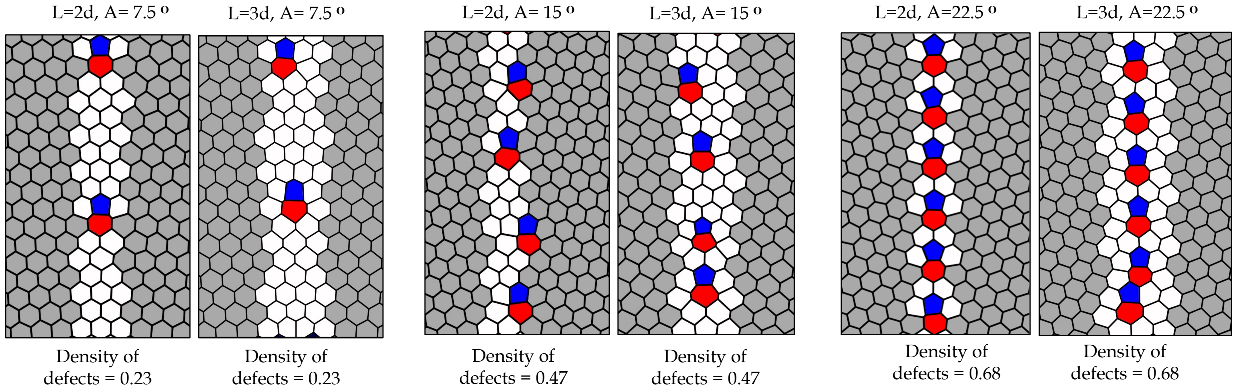


Fig. S6: Samples from the experimental data. Each pair shows similar patterns of the cells built within the gaps for the same angle of misalignment ($A$), but different values for the horizontal shift ($L$). The density of defects for each pair stays constant, even though the horizontal distance increases.

#### 1.4 Incomplete Trials

Fig. S7, and Fig. S8 show instances of a few experimental frames that were not included in our analysis. Bees did not merge the two sides of the gap in both images shown in Fig. S7 during the time frame of our experiments. This is probably due to the large horizontal distance between the given hexagonal panels. We excluded trials where bees constructed drone cells, due to the poly-crystal nature of the lattice. For example, Fig. S8 shows a scenario with multi drone cells. The cells built in the gap are shown in Fig. S8 B with blue, white, and red for 5-, 6-, and 7-sided cells respectively, and are generally larger than the cells that are built on the given patterns—see the cell size distribution in Fig. S8 C. Fig. S8 D shows how bees make use of the larger cells in the gap by turning those into drone comb which is approximately 1.5X larger than the worker comb.

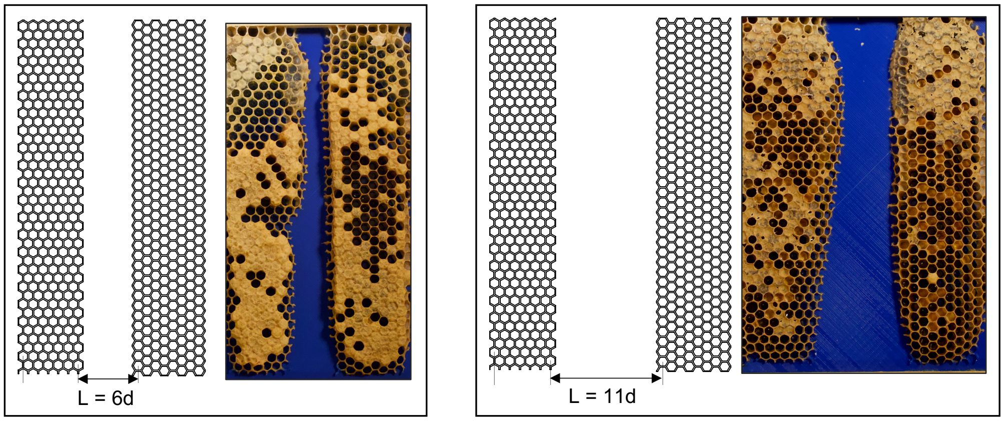


Fig. S7: Example of a couple of experimental frames where the two sides of the comb did not merge during the time frame of our experiments.

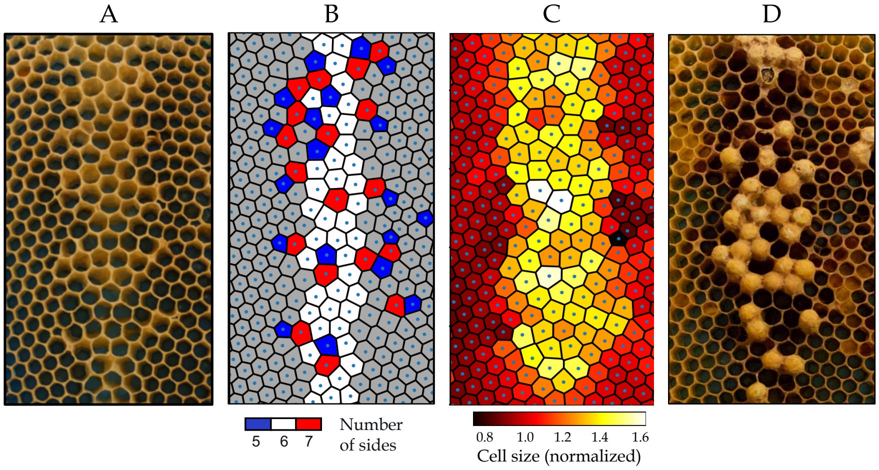


Fig. S8: A) An instance of comb construction on one of our experimental frames with $A = 15^\circ$, $L = 3d$, and $h = 0$. B) Voronoi reconstruction of the comb in panel A with cells colored by the number of sides shows a non-regular pattern of defects which is not the same as the patterns we usually observed for this parameter combination. C) Voronoi reconstruction colored by the area of the cells shows that larger defective cells are built in the gap whereas the surrounding hexagonal cells are generally smaller. D) The same crop shown in panel A photographed after a month shows that the bees make use of the large cells in the gap by turning them into drone comb, which typically consists of larger cells to grow drones.

#### 1.5 Number of cells in the gap

To obtain a correct estimate of the number of moving cells in our model, we counted the number of cells that are built within the gap for a fixed crop size by overlaying the experimental results on the design sheet, see an example in Fig. S9 B. Table A shows those numbers for all the configurations.

A

| Angle | Distance | Number of cells in the gap |
|---|---|---|
| **0** | **2d** | 61 |
| **7.5** | **2d** | 48 |
| **15** | **2d** | 49 |
| **22.5** | **2d** | 49 |
| **30** | **2d** | 30 |
| **0** | **3d** | 86 |
| **7.5** | **3d** | 73 |
| **15** | **3d** | 74 |
| **22.5** | **3d** | 77 |
| **30** | **3d** | 59 |
| **0** | **4d** | N/A |
| **7.5** | **4d** | 96 |
| **15** | **4d** | 97 |
| **22.5** | **4d** | 99 |
| **30** | **4d** | 84 |

B

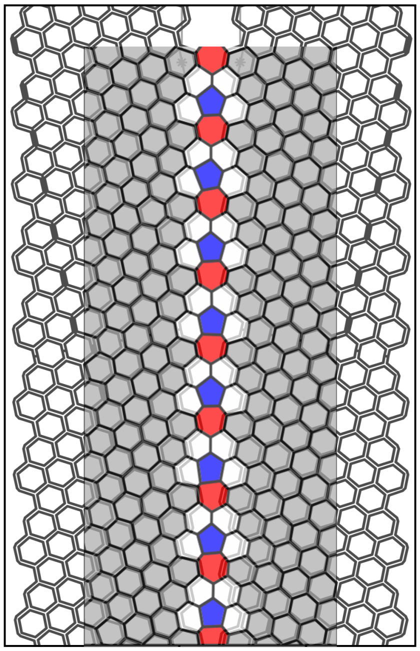

Fig. S9: A) The average number of cells built in the gap for each experiment configuration. B) An overlay of one of our experimental crops on the design sheet for $A = 22.5°$, and $L = 2d$, which is also highlighted in Table A. The gray-shaded hexagons are the cells that are built on the given foundation. Other cells are built in the transitional area. The number of transitional cells is equal to 49.
